# Stochastic resonance is behind single cell proteomics success

**DOI:** 10.1101/2025.08.22.671757

**Authors:** Zhaowei Meng, Roman A. Zubarev

## Abstract

Stochastic resonance is a phenomenon of noise-aided detection of below-threshold signal at the expense of the linearity of signal quantification. Here we demonstrate that peptide signal from single cell proteomes multiplexed by tandem mass tag (TMT) with a carrier proteome is below the nominal detection threshold in the Orbitrap mass analyzer and is registered in the mass spectra largely due to stochastic resonance amplification. This amplification is of the order of 0.5-2.5 single cell equivalents. Consequently, spiking by the corresponding amount of a constant “pedestal” proteome lifts the signal of the analyte single cell proteome above the detection threshold, improving detection and largely restoring linearity of quantification.

## INTRODUCTION

Stochastic resonance (SR) is a phenomenon in which makes detectable a below-threshold signal by, paradoxically, adding noise to it^1, 2^. The omnipresent “endogenous” noise also helps detection of sub-threshold signals. The enhancement in signal detection due to SR can be as large as 1 to 2 orders of magnitude^2^, but it often comes at the expense of the signal linearity, affecting quantification accuracy. SR has long ago found practical applications in different areas of analytical sciences, including mass spectrometry (MS), particularly in chromatography-coupled MS analyses (GC-MS and LC-MS). For instance, Xiang et al. applied periodic modulation to the LC-MS detection of granisetron in plasma, and reduced using an SR-based algorithm the limits of detection and quantification fivefold and tenfold, respectively^3^. Similarly, Zhang et al. SR-detected weak chromatographic signal of glyburide in plasma, significantly boosting the signal-to-noise ratio of low-level peaks^4^. The same team applied SR to the analysis of roxithromycin in beagle dog plasma^5^. Deng et al. used SR in detection of multiple weak peaks in ultra-performance LC coupled to time-of-flight (TOF) MS^6^. These advances suggest that stochastic resonance may offer a valuable tool for enhancing sensitivity in MS workflows, especially where traditional hardware or separation improvements have reached practical limits. However, most of SR-related studies were performed before Fourier transform (FT) mass analysers, such Orbitraps, became commercially available and there seems to be no published examples of SR use in FT MS.

When first reports on the modern single cell proteomics (SCP) appeared, they were met with skepticism in the proteomics community, which delayed the first SCP publication^7^. The skepticism was based on several studies in which serial dilution of the bulk proteome failed to produce useable signal below the equivalent of 5-10 cells. For instance, Bruderer et al. performed in 2015 serial dilutions using a human cell-derived proteome standard analyzed with state-of-the-art data-independent acquisition (DIA/SWATH-MS)^8^. In their analysis of fourfold dilution series progressing into the lower sample amounts, signals corresponding to the proteome content of fewer than 10 cells fell below practical limits of detection, yielding insufficient peptide identifications and quantitative data for robust analysis. This finding reflected a broader trend observed in earlier bulk proteome experiments, where random protein sampling in data-dependent acquisition (DDA), ion suppression, and limited instrument sensitivity collectively resulted in a rapid drop-off of proteome coverage at sub-nanogram sample amounts, while only 150-200 pg of protein can be extracted from a single cell^9^. Yet Single Cell ProtEomics by Mass Spectrometry (SCoPE-MS) by the Budnik et al. ^7^provided definite detection and quantification of hundreds and eventually thousands of proteins from single cells, gradually winning over the skepticism of critics. The “secret” of SCoPE MS was the use of the “carrier proteome” composed of 150-250 cells multiplexed by tandem mass tag (TMT) with a multitude of single cell proteomes. The signal from the carrier proteome is an order of magnitude above the conventionally accepted detection limit in the Orbitrap analyzer, ensuring detection of thousands of peptides, while the TMT reporter ions provided information on the peptide abundance in single cells. This experimental scheme allowed for easy verification in most Orbitrap-equipped proteomics laboratories, and SCoPE MS quickly became confirmed valid. What was however rarely discussed is that the average signal of the reporter ions from single cell proteomes was not 150-250 times smaller than that of the carrier proteome, as expected, but often at order of magnitude higher than that^10^. A similar effect was also observed when SCoPE MS was applied to single bacterium proteome^11^. While multiple artifacts could possibly contribute to this phenomenon, we postulate here that the main contributor to signal enhancement in SCoPE MS is stochastic resonance. Here we exploit this issue by systematic experiments.

## RESULTS

To verify whether SR plays a role in SCoPE MS signal detection, we extracted bulk proteome from A549 human cells, digested it as customary in proteomics (see Methods), labelled with TMT16 and then we performed serial dilution. Assuming single cell proteome weight to be 0.2 ng, in the rest of the discussion we will use single cell equivalents. In the TMT labelling scheme (**Table 1**), together with the carrier proteome of 200 cells the proteomes of 1 to 20 cells were multiplexed in two replicates each. To avoid carry-over from the carrier proteome channel, the corresponding TMT channel remained empty, as conventional in SCoPE MS^7^.

**Table 1.**
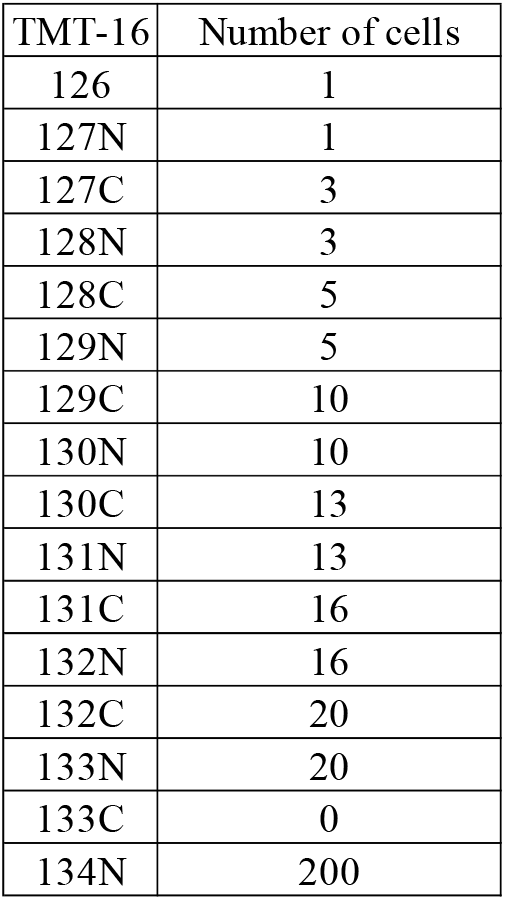
TMT labelling scheme for detecting the SR amplification.

| TMT-16 | Number of cells |
| --- | --- |
| 126 | 1 |
| 127N | 1 |
| 127C | 3 |
| 128N | 3 |
| 128C | 5 |
| 129N | 5 |
| 129C | 10 |
| 130N | 10 |
| 130C | 13 |
| 131N | 13 |
| 131C | 16 |
| 132N | 16 |
| 132C | 20 |
| 133N | 20 |
| 133C | 0 |
| 134N | 200 |

After LC-MS/MS analysis the abundances of each of 659 quantified proteins were plotted against the number of cells in the sample, from n to N cells. The negative ratio of the intercept and slope of a linear regression for given n and N provided the detection threshold (limit) as the number of cells at which the abundance of a given protein becomes equal to zero. The null hypothesis assumes no SR effect, in which case the average threshold would not change significantly with n and N. However, **Figure 1** clearly shows that the threshold becomes significantly lower for smaller n and N, from 2.5 cells for n=10 and N=200 cells to 0.5 cells for n=1 and N=5 cells, consistent with SR-increased signal at low cell load. But the latter threshold value means that even for a few-cell load the SR effect was not sufficient to reduce the detection threshold to zero. Thus up to a half (by abundance) of the single cell proteome signal remains undetected at the SCoPE MS conditions or consist of pure noise. Indeed, in 40.2 % of the cases the TMT channels corresponding to single cells were found empty, as opposed to 13.5 % of the cases for the five-cell equivalents. In SCP, missing values represent a big issue, most often solved by imputation of an arbitrary value^12^, which hampers quantification.

**Figure 1.**
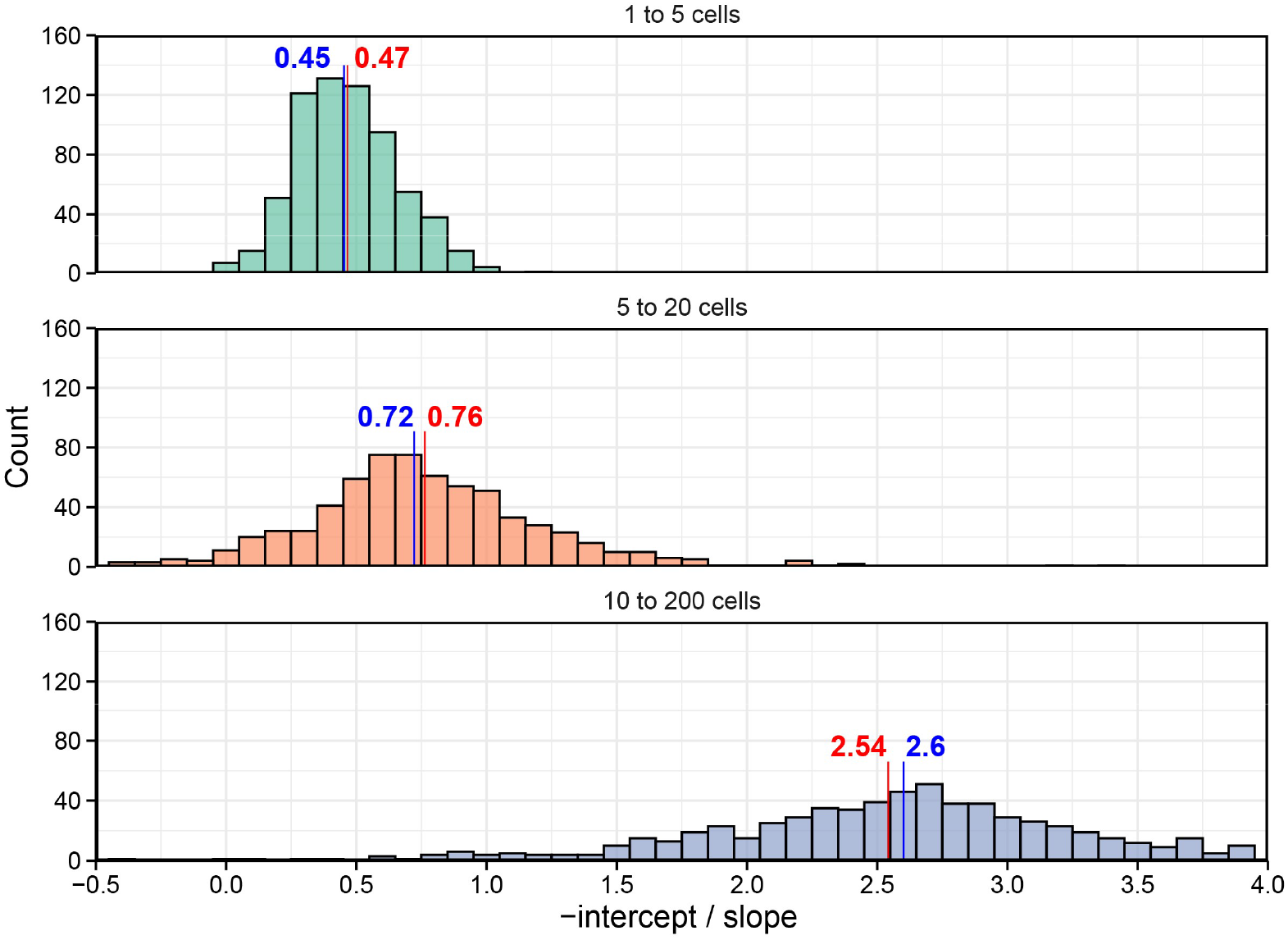
The distribution of the detection threshold (limit) for different intervals of cell numbers. The intercept and slope were derived from the linear regression between protein abundances and the number of cells, from n to N cells. The negative of intercept-to-slope ratio provided the number of cells where the abundance of a given protein becomes zero, revealing the detection threshold (limit). The median (blue) and average (red) values are shown.

We therefore supposed that spiking the samples with a constant “pedestal” background of a single cell equivalent will reduce the SR-enhanced signal threshold to zero or below and improve the SCoPE MS performance, increasing the number of quantified peptides and proteins and reducing the number of missing values. Moreover, we expected that the reduction of missing values or the channels filled with noise will improve the replicate-to-replicate coefficient of variability (CV).

For the proof of principle experiment we created another TMT set (**Table 2**) containing 200 cells as a carrier proteome and from n=0 to N=2.5 cell equivalents to which one cellular proteome was always added. The LC-MS/MS results confirmed our expectations (**Figure 2A**): for n=0 and N=2.5, the detection threshold was negative both on average (-0.61 cells) as well as as median (-0.62 cells) for all detected proteins. Even for the case of n=1 and N=2.5, the average detection threshold was still negative (-0.57 cells), the same as for 96.8 % of all detected proteins (**Figure 2B**). Moreover, the replicate-to-replicate CV values diminished from the median 21% without the add-on to 16% with a one-cell “pedestal” background spiked in (**Figure 2C**).

**Table 2.**
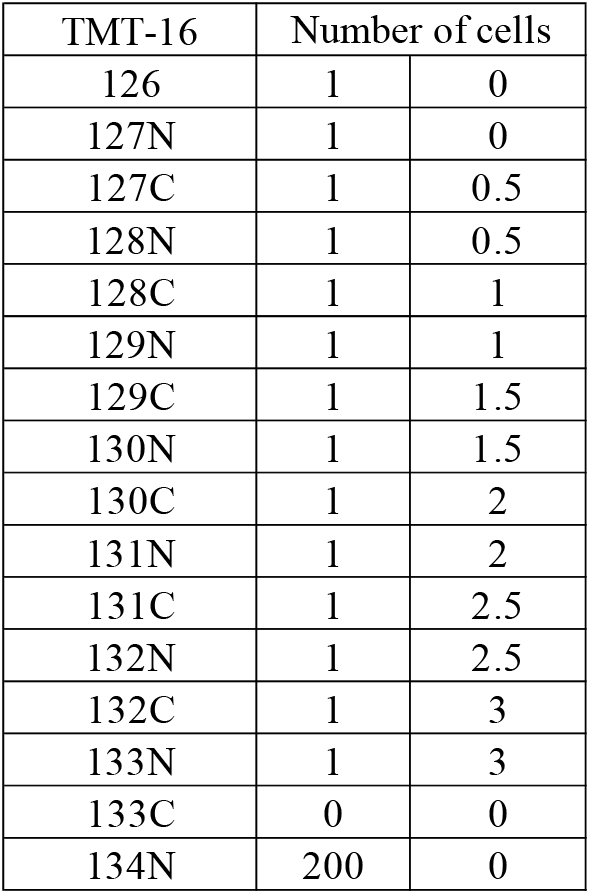
TMT scheme for the proof of principle experiment with a constant add-on spiked into SCP samples.

| TMT-16 | Number of cells |  |
| --- | --- | --- |
| 126 | 1 | 0 |
| 127N | 1 | 0 |
| 127C | 1 | 0.5 |
| 128N | 1 | 0.5 |
| 128C | 1 | 1 |
| 129N | 1 | 1 |
| 129C | 1 | 1.5 |
| 130N | 1 | 1.5 |
| 130C | 1 | 2 |
| 131N | 1 | 2 |
| 131C | 1 | 2.5 |
| 132N | 1 | 2.5 |
| 132C | 1 | 3 |
| 133N | 1 | 3 |
| 133C | 0 | 0 |
| 134N | 200 | 0 |

**Figure 2.**
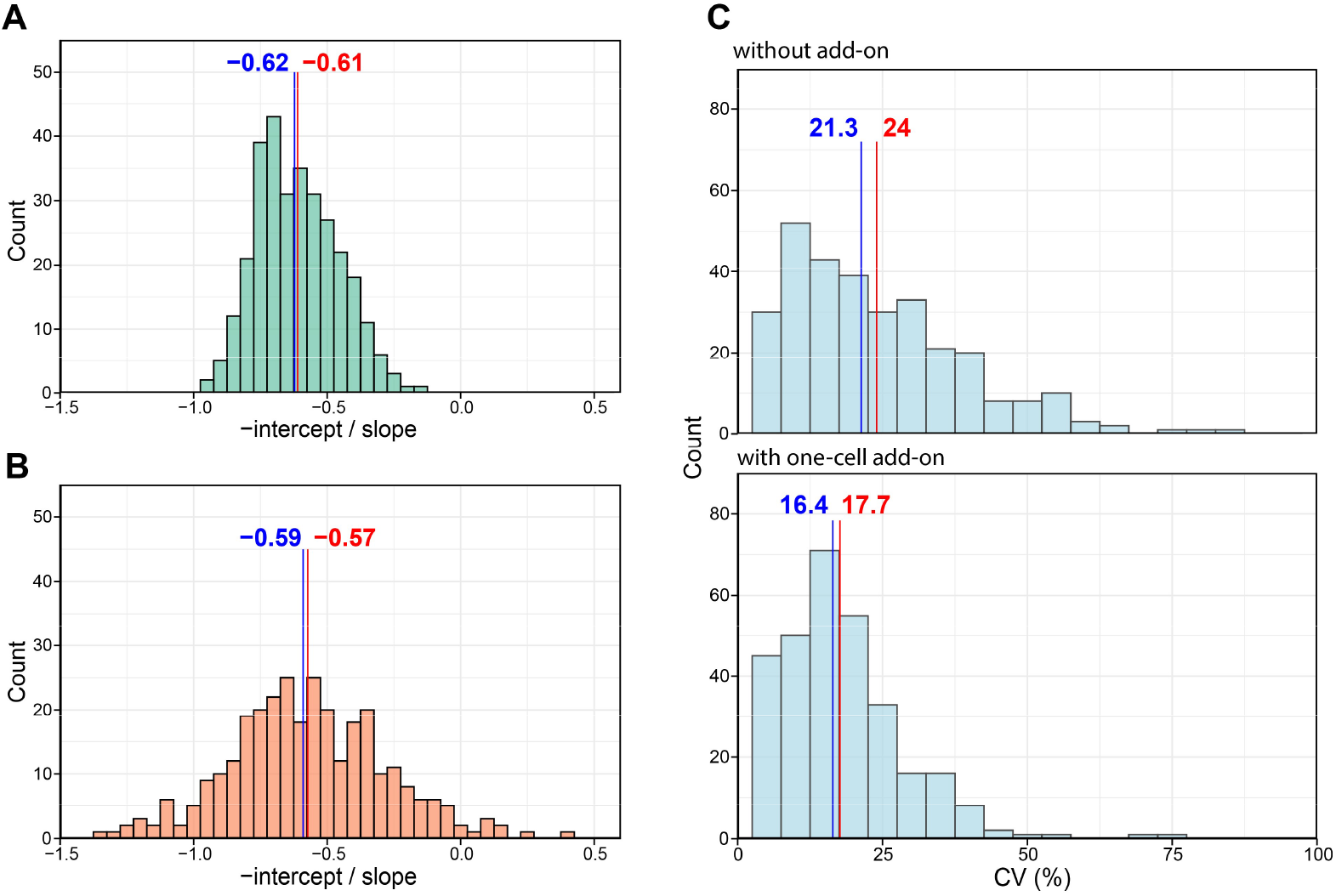
The proof of principle experiment with a constant add-on cellular proteome. (A-B) The distribution of the detection threshold for n to N cell equivalents to which one cellular proteome was always added, (A) n=0 and N= 2.5, (B) n=1 and N= 2.5. (C) The replicate-to-replicate CV values without the add-on and with a one-cell “pedestal” background spiked in. The median (blue) and average (red) values were labelled.

## DISCUSSION

Our results confirmed that SR is the likely reason for the detection limit decrease in the Orbitrap Lumos mass spectrometer from a ≈2.5 cell equivalent for a ≥10 cell load to a ≈0.5 cell equivalent in single cell analysis. The natural detection limit of ≈2.5 cells is broadly consistent with the earlier studies with less sensitive mass spectrometers that concluded the necessity of at least nanogram sample load^8, 9^. The 5-fold enhancement in the detection threshold is also in line with the known SR ability to increase the signal 1-2 orders of magnitude ^2^ and the previous use of SR in mass spectrometry^3-6^. Despite this enhancement, about half (by abundance) of SCP signal is below the detection threshold in a typical SCoPE MS setting, with the missing values filled either by noise or by arbitrary value imputation during bioinformatic analysis^12^. Even though in both cases the quantification aspect is affected, data quality may remain sufficient for analyses heavily relying on the protein abundance changes^10^. Despite that, every increase in data quality, such as reduction in missing values and in CVs, is welcome in such a rapidly developing area as single cell proteomics. Here we demonstrated that addition of a constant pedestal (background proteome) equivalent to a single cell lifts all SCP signals above the detection threshold in an advanced Orbitrap mass spectrometer and enables linear quantification of even low-abundant proteins.

## CONCLUSIONS

Regrettably, the ‘number game’ that plagued the proteomics field before it reached maturity ^13^ seems to have become fashionable once again, this time in single cell proteomics. When this immature phase is over, the focus of SCP research will invariably shift to quantification. Once again, lower CVs will become more important than elevated numbers of detected proteins^14^, and the “one hit wonders” of which significant part of contemporary SCP data consists of will once more be deemed unsuitable for quantification^15^. When serious applications will demand reliable quantitative performance, the approach presented here may become useful in single cell proteomics workflows.

## Acknowledgements

This work was supported by grants from VR 2021-05223, Cancerfonden 22 1967 Pj and EU ARIADNE VIBE consortium.

## Author contributions

R.Z. conceived the project and supervised the study. R.Z. and Z.M. designed the experiments. Z.M. performed the experiments and analyzed the data. R.Z. and Z.M. wrote the manuscript. All authors approved the final version of the manuscript.

## Methods

### Cell Culture

Human lung carcinoma A549 cells (ATCC) were grown at 37°C in 5% CO_2_ using Dulbecco’s Modified Eagle Medium (Lonza) supplemented with 10% FBS superior (Biochrom) and 100 units/mL penicillin/streptomycin (Gibco). Low-number passages (<10) were used for the experiments. Cellular lysates were obtained in PBS by five cycles of freezing cells in liquid N_2_ and then thawing at 25 °C, then cleared by centrifugation at 14,000 rpm for 10 min at 4 °C. The protein concentration in the lysate was measured using Pierce BCA kit (Thermo Fisher).

### Reduction, Alkylation and Digestion

500 µg of A549 lysate was transferred to a Lobind tube. Dithiothreitol (DTT) was added to a final concentration of 10 mM and samples were incubated for 30 min at 55 °C. Subsequently, iodoacetamide (IAA) was added to a final concentration of 30 mM and samples were incubated at 25 °C for 30 min in the darkness. Proteins were precipitated using methanol and chloroform. The precipitated protein was collected by centrifugation (17,000 × g, 10 min), and the supernatant was removed. The protein pellet was air-dried for 3 min, then dissolved in 150 µL of 20 mM EPPS buffer at pH 8.2 containing 8 M urea for 10 min. 20 mM EPPS buffer (pH 8.2) was added to dilute urea to 4 M. Then Lysyl endopeptidase (LysC) (Wako) was added at a 1:50 w/w ratio, the samples were incubated at 30 °C for 16 h. The samples were diluted with 20 mM EPPS to a final urea concentration of 1 M, and trypsin was added at a 1:50 w/w ratio, followed by incubation at 37 °C for 6 h. The resulting peptides sample was acidified by TFA to pH 1-2, cleaned using Sep-Pak cartridges (Waters), and dried using DNA 120 SpeedVac Concentrator (Thermo Fisher).

### TMTpro Labeling

After drying the peptides were resuspended in 20 mM EPPS buffer at pH 8.2. The pepide concentration was measured using Pierce BCA kit (Thermo Fisher). 25 µg of peptides was transferred to each Lobind tube (16 tubes in total). TMTpro 16plex reagents dissolved in acetonitrile (ACN) were added 5× by weight to each sample with a final ACN concentration of 30%, followed by incubation for 2 h at 25 °C. The reaction was quenched by adding 1.5% hydroxylamine.

### Amplification Factor Determination

130 µL of 10 ng/µL TMTpro-labeled peptides solution was prepared for each channel except for 133C (empty) and 134N (carrier proteome). 50 µL of 17.5 ng/µL TMTpro 134N-labeled peptides solution was prepared. The single cell proteome weight was considered to be 0.2 ng. 10 ng/µL TMTpro-labeled peptides for the channels from 126 to 133N were combined using the volume in Table 1 to generate 272 µL of combined sample, 16.3 µL of which was mixed with 13.7 µL of 17.5 ng/µL carrier proteome (channel 134N) for the final sample solution. The injection volume was 5 µL for LC-MS/MS analysis.

### Add-On Level Determination

10ng/µL TMTpro-labeled peptides for the channels from 126 to 133N were combined using the volume in Table 2 to generate 70 µL of combined sample, 4.2 µL of which was mixed with 13.7 µL of 17.5 ng/µL carrier proteome (channel 134N) and 12.1 µL of 0.1% TFA-2% ACN in water for the final sample solution. The injection volume was 5 µL for LC-MS/MS analysis.

### LC-MS/MS Analysis

The samples were analyzed by LC-MS/MS using a Orbitrap Fusion Lumos mass spectrometer equipped with an EASY Spray Source and connected to an Ultimate 3000 RSLC nano UPLC system (all - Thermo Fisher). Injected sample were preconcentrated and further desalted online using a PepMap C18 nano trap column (2 cm X 75 μm; particle size, 3 μm; pore size, 100 Å; Thermo Fisher) with a flow rate of 3.5 μL/min for 6 min. Peptide separation was performed using an EASY-Spray C18 reversed-phase nano LC column (Acclaim PepMap RSLC; 50 cm X 75 μm; particle size, 2 μm; pore size, 100 Å; Thermo Fisher) at 55°C and a flow rate of 300 nL/min. Peptides were separated using a binary solvent system consisting of 0.1% (v/v) formic acid (FA), 2% (v/v) acetonitrile (ACN) (solvent A) and 98% ACN (v/v), 0.1% (v/v) FA (solvent B) with a gradient of 3−17% B in 56 min, 17−25% B in 21 min, 25−35% B in 13 min, 35−85% B in 4 min. Subsequently, the analytical column was washed with 85% B for 8 min before re-equilibration with 3% B.

To identify and quantify TMT-labelled peptides, we utilized SPS MS3 methods. In the MS2 approach, a full MS (MS1) spectrum was first acquired in the Orbitrap analyzer with mass-to-charge ratio (m/z) range from 350 to 1500, nominal resolution 120,000, automated gain control (AGC) target 1 × 10^6^, and maximum injection time of 100 ms. The most abundant peptide ions were automatically selected for subsequent MS/MS (MS2) analysis with a minimum intensity threshold of 2 × 10^4^ and a 45 s dynamic exclusion time. MS2 spectra were acquired in the Orbitrap analyzer with the following settings: quadrupole isolation window 0.7 Th, AGC target 2 × 10^4^, maximum injection time 250 ms, fragmentation type HCD, normalized collision energy 38%, nominal resolution 60,000, fixed first m/z 100. The number of MS2 spectra acquired per each MS1 spectrum was determined by setting the maximum cycle time for MS1 and MS2 spectra to 2 s (using a top speed mode).

### Data Processing

The raw LC-MS/MS data were analysed by MaxQuant, version 2.2.0.0. The Andromeda search engine was employed to perform MS/MS data matching against the UniProt Human proteome database (version UP000005640_9606, 20607 human sequences). Enzyme specificity was trypsin, with maximum two missed cleavages permitted. When needed, semi-tryptic option was specified for the identification of semi-tryptic peptides in the MS2 dataset. Cysteine carbamidomethylation was set as a fixed modification, while methionine oxidation, N-terminal acetylation, asparagine or glutamine deamidation were used as a variable modification. 1% false discovery rate was used as a filter at both protein and peptide levels. Default settings were employed for all other parameters. Peptide quantification was executed using TMTpro 16plex.

